# Neur-Ally: A deep learning model for regulatory variant prediction based on genomic and epigenomic features in brain and its validation in certain neurological disorders

**DOI:** 10.1101/2025.01.27.635013

**Authors:** Anil Prakash, Moinak Banerjee

**Affiliations:** Human Molecular Genetics Lab, Neurobiology and Genetics Division, Rajiv Gandhi Centre for Biotechnology, Thiruvananthapuram, Kerala, 695014, India; Department of Biotechnology, University of Kerala, Kariavattom, Thiruvananthapuram, Kerala, India

**Author notes:** To whom correspondence should be addressed. **<u>Corresponding author,</u>** Moinak Banerjee PhD, Human Molecular Genetics Laboratory, Rajiv Gandhi Centre of Biotechnology [RGCB], Thycaud Post, Poojappura, Thiruvananthapuram, Kerala 695014. India.

**Keywords:** SNPs, Non-coding, Epigenetics, regulatory, chromatin accessibility, histone modifications, transcription-factor [TF] binding, Brain, Neurological disorders

## Abstract

Large scale quantitative studies have identified significant genetic associations for various neurological disorders. Expression quantitative trait loci [eQTL] studies have shown the effect of single nucleotide polymorphisms [SNPs] on the differential expression of genes in brain tissues. However, a large majority of the associations are contributed by SNPs in the noncoding regions which can have significant regulatory function but are often ignored. Besides mutations that are in high linkage disequilibrium [LD] with actual regulatory SNPs will also show significant associations. Therefore, it is important to differentiate a regulatory non-coding SNPs with a non-regulatory one. To resolve this, we developed a deep-learning model named Neur-Ally, which was trained on epigenomic datasets from nervous tissue and cell line samples. The model predicts differential occurrence of regulatory features like chromatin accessibility, histone modifications and transcription-factor [TF] binding on genomic regions using DNA sequence as input. The model was used to predict the regulatory effect of neurological condition specific non-coding SNPs using in-silico mutagenesis. The effect of associated SNPs reported in Genome-wide association studies [GWAS] of neurological condition, Brain eQTLs, Autism Spectrum Disorder [ASD] and reported probable regulatory SNPs in neurological conditions were predicted by Neur-Ally.

## INTRODUCTION

Understanding the genetics of complex disorders are extremely difficult due to the heterogeneity within and between populations [1]. The decreasing cost of sequencing and the emergence of custom genotyping arrays have resulted in an increase of quantitative genetic studies like GWAS and eQTL analysis. Linkage disequilibrium between regulatory and normal SNPs can lead to the emergence of non-causative genetic variants in association results [2]. Identifying the actual causative variant using experimental techniques will be extremely difficult. The common significant variant may be having a narrow effect on the phenotype and the combined effect of multiple variants will be needed for the phenotype to occur [2]. So functional screening of a large number of associated single genetic variants will be challenging.

Non-coding genetic variants are highly enriched in risk variants, identified by quantitative genetic analysis of complex diseases [3]. The effect of such mutations can be indirect and regulatory in function. The effect of coding mutations like missense and nonsense SNPs can be studied by analyzing the structural changes introduced in the normal protein. In contrast, the regulatory effect of a non-coding SNP will be difficult to decipher. The regulatory landscape can be cell line or tissue specific and this adds to the problem. Large number of epigenomic datasets from different tissues and cell lines are available from the Encode project [4]. The incorporation of regulatory features improved the modelling and representation of complex diseases [5]. Hence the use of tissue specific regulatory datasets will aid in understanding more about the interplay between the regulome and complex diseases [6].

Through the use of ChIP-seq, researchers have made substantial progress in locating transcription factor binding sites, investigating the regulatory roles of transcription factors in gene expression, and mapping histone modifications throughout the genome [7]. ChIP-seq analysis has increasingly been combined with other functional genomics techniques, facilitating a deeper understanding of the mechanisms that regulate gene expression [8]. A wide array of quantitative methodologies has significantly contributed to the progress in assessing chromatin accessibility. Continuous improvements in integrated analysis are expected to enhance our comprehension of the intricate relationships among DNA accessibility, gene expression, genetic variants, protein interactions, transcription and subsequent phenotypes [9].

The regulation of genes, which includes transcription and alternative splicing, is fundamentally influenced by DNA- and RNA-binding proteins. Recent advancements in deep learning methodologies have facilitated the prediction of the sequence specificities of these proteins, thereby enhancing our understanding of regulatory mechanisms [10]. Deep learning models are capable of discerning regulatory sequence patterns from extensive regulatory datasets, such as chromatin accessibility data. This capability allows for the prediction of chromatin effects resulting from sequence modifications with single-nucleotide precision. As a result, these models have significantly improved the prioritization of functional variants, which encompasses eQTLs and variants linked to diseases [11].

Computational tools that are created using cell or tissue specific datasets will help in better understanding of the diseases that are connected to those types of tissues or cells. This prompted us to create a model for all neurological conditions, which can be trained on neuronal specific epigenomic datasets and that will subsequently help in the variant effect prediction of mutations specific to neurological conditions. Several deep learning models have been developed with applications in biology [12]. They will learn the linear and non-linear relationships within the vast amount of data that is available. Attention layers have improved the performance of various Natural language processing [NLP] tasks which in turn helped in the analysis of sequence datasets [13]. With this in mind, we developed a deep learning model called Neur-Ally. The model has convolution and attention layers incorporated into the architecture.

The model after training can be used to predict the regulatory effect of SNPs specific to neurological conditions. These SNPs can be chosen from significant GWAS associations or candidate genetic association studies for neurologic conditions [14]. Alternatively, eQTL SNPs which regulate the expression of genes can also be used for the prediction [15]. In case of variants occurring in non-neuronal tissues contributing to the phenotype, epigenomic datasets from those samples can also be processed using the data processing codes available with the model. For neurological condition specific mutations, the pre-trained weights along with the codes are available for testing. In short, Neur-Ally will help in identifying the regulatory potential of SNPs in the brain, based on the differential epigenomic signatures in response to in silico mutagenesis.

## MATERIALS AND METHODS

### Model Architecture

The genomic bins where epigenomic labels overlap are used as input to the model. The 200base pair [bp] sequence of the bin along with the flanking regions [1800 bp] are subjected to vectorization, word embedding and positional encoding to create a multi-dimensional tensor. Then, it is fed into subsequent layers of 1D convolution and max pooling twice, followed by Multi-Head Attention layers. The output from the final attention layer is fed into the Keras Dense layer [16], this is followed by reduction in the dimension of the tensor using squeeze operation. The final Dense layer and sigmoid activation function, provides the output of the probability of regulatory signatures in the genomic bin region used as the input. The flow diagram of the model architecture is shown in Figure 1.

**Figure 1:**
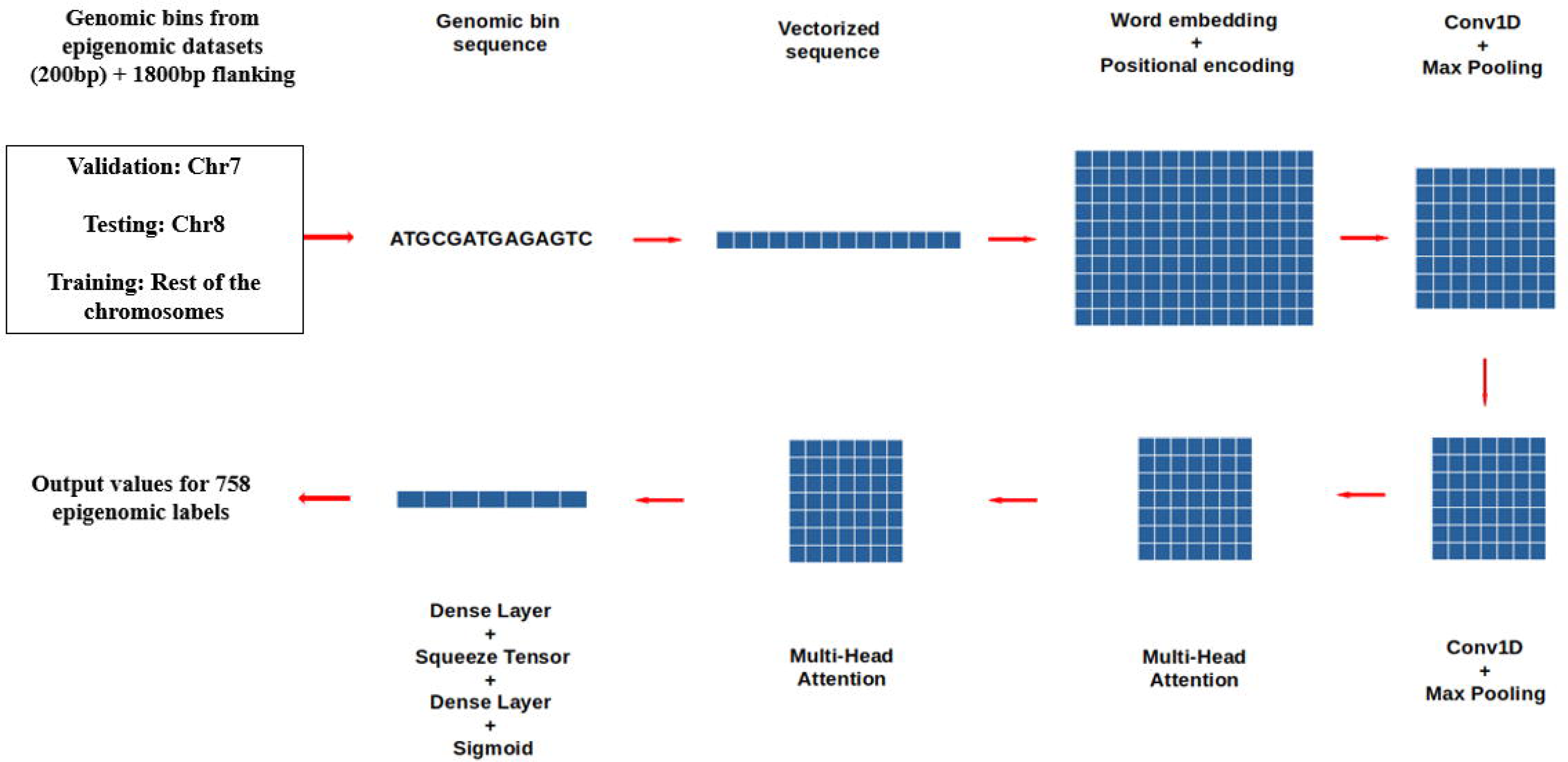
Neur-Ally architecture. Flowchart diagram of the model architecture.

### Data processing

Epigenomic datasets regarding chromatin accessibility [ATAC-Seq, DNase-Seq], Histone modifications and TF binding pertaining to tissue type or cell type were selected for processing the data [Figure 2]. The narrow peak bed files of nervous tissue and cell samples were extracted from the Encode Project. Genomic bins of 200 bp length were selected as positive samples if the epigenomic signature is overlapping more than half of it. Genomic bins with low mappability were excluded from the analysis. The processed dataset was split into training, validation and testing based on the chromosome number of the genomic bin. Those belonging to chromosome 7 and chromosome 8 were used for validation and testing, whereas the remaining ones were kept for training. For testing the model predictions, Area Under the Curve of Receiver Operator Characteristic [AUROC] and Precision Recall [PR-AUC] curves were estimated.

**Figure 2:**
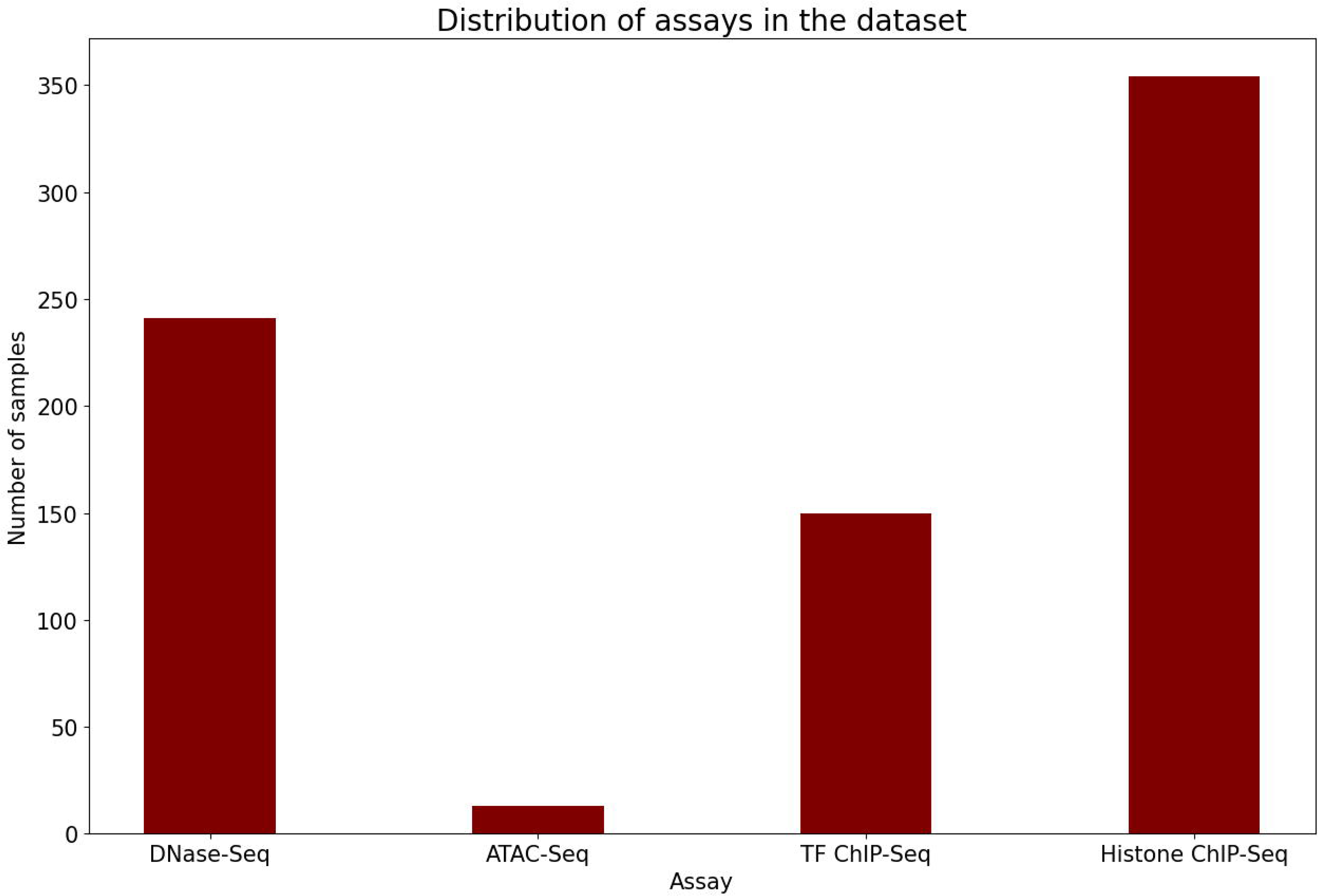
Assay distribution. Bar graph of assays used for data preprocessing.

### Variant effect prediction

As the model was trained on genomic sequence and regulatory labels, it can learn the contribution of sequence features to the prediction. Thus, the prediction of regulatory labels upon giving altered sequences as input can shed light on the regulatory effect of mutations. Therefore, we predicted the regulatory effect of SNPs specific to neurological conditions [SNMs] by in silico mutagenesis. The sequence of the genomic bin harboring the mutation was extracted from the reference genome. Another input sequence was generated by altering the nucleotide at the mutation site. Both the inputs were given to the model in a sequential manner. The predictions of the regulatory labels were compared for both the sequences to estimate the SNP Activity Difference [SAD] score [17].

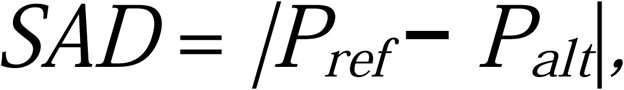

where *P_ref_* is the probability of regulatory labels predicted on the reference sequence, whereas *P_alt_* is the probability of regulatory labels predicted on the mutated sequence. SAD score is the absolute difference between *P_ref_* and *P_alt_*.

To identify significant regulatory variants, we created a negative non-regulatory set of SNPs from the 1000 genome dataset [18]. A million variants were randomly selected and a negative set of SNPs were created by filtering GWAS and eQTL variants and those occurring in exonic or candidate cis-regulatory regions. The significance of the regulatory effect of the predicted variants were estimated using the E-value method [19]. E-value of an epigenomic target for a particular SNP is defined as the ratio of SNPs from the negative non-regulatory set having higher SAD score for the same target. The same number of positive and negative variant sets are used for E-value prediction. The selection of negative samples is repeated ten times and the mean E-value is selected for comparison. SNPs with an E-value of “1e-05” or less are considered as significant.

Neur-ally was trained on epigenomic datasets with coordinates according to the hg38 human reference build. So, the input variant coordinates have to be based on the hg38 build. While using SNP datasets belonging to older genomic builds, the coordinates were converted to the latest genomic build using liftover tools. Chromosomal coordinates belonging to conversion unstable positions were excluded from the analysis [20].

### Model prediction on GWAS of neurological conditions and eQTL SNPs

The trained model was used to identify the differential regulatory label prediction of neurological condition specific SNPs extracted from the GWAS catalog and eQTL variants in the brain tissues. For the neurological condition specific GWAS SNPs, “GWAS catalog v1.0” dataset was extracted and associated SNPs were selected based on matching keywords in the disease or trait column. The following keywords were used for the filtering: “alzheimer”, “epilepsy”, “multiple sclerosis”, “parkinson”, “autism”, “attention deficit”, “schizophrenia”, “bipolar”, “major depressive”. Significant variant-gene pair datasets of the neuronal tissues from the GTEx portal were used for selecting the neurological condition specific eQTLs. Top 1000 significant eQTL SNPs from each sample were used for creating the list of eQTLs to be tested.

### Model prediction on ASD GWAS and brain regulatory SNPs

The E-value threshold of “1e-05” is a stringent one, but since the calculated E-value will depend on the number of variants present in the positive set, so we tried to restrict the prediction to Autism Spectrum Disorder associated SNPs from the GWAS catalog. Hence, we used the keywords, “Asperger disorder”, “Autism”, “Autism spectrum disorder”, to select the variants from the dataset. Next, we wanted to test the model performance on reported probable brain regulatory SNPs [21]. As the number of positive variants are few, 200 negative variants were sub-sampled for comparing the SAD scores.

## RESULTS

### Model performance

The Binary Cross-Entropy loss values and metrics during training and validation over 39 epochs are depicted in Figure 3. The metric values generated by Keras were approximated ones and the individual metric values of each epigenomic label were generated using scikit-learn [22] [Supplementary Table S1]. The prediction of chromatin accessibility assay labels had a mean AUROC of 0.93 and PR-AUC of 0.23 [baseline PR-AUC is 0.01]. Histone modifications had a mean AUROC of 0.84 and PR-AUC of 0.29 [baseline PR-AUC is 0.03]. Transcription factor binding labels had a mean AUROC of 0.87 and PR-AUC of 0.22 [baseline PR-AUC is 0.01].

**Figure 3:**
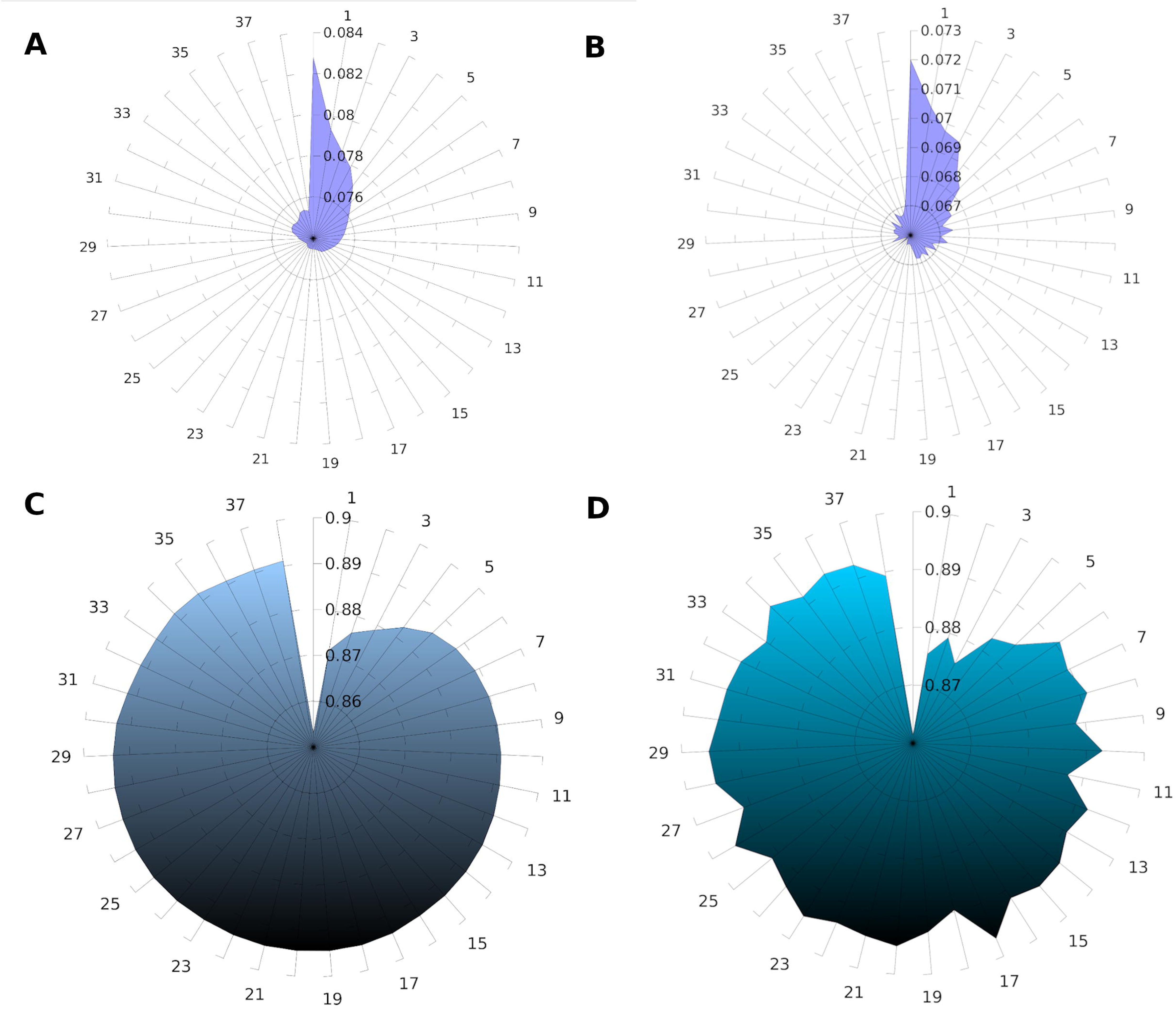
Training and validation metrics. Radar plots showing A.Training loss, B. Validation loss, C. Training mean AUROC, D. Validation mean AUROC over subsequent epochs. Epochs are depicted in circular axes and metric or loss values in radial axes.

### GWAS and neurological condition specific eQTL variants

GWAS associated SNPs were selected after removing GWAS Catalog variants occurring in the coding regions. 7663 neurological condition specific SNPs were extracted using keywords and selected as the positive variant set. 48 SNPs were showing significant E-values after comparing with the negative variant set at a threshold of “1e-05” [Figure 4A, Supplementary Table S2]. Using a less stringent threshold can reveal more possible regulatory variants.

**Figure 4:**
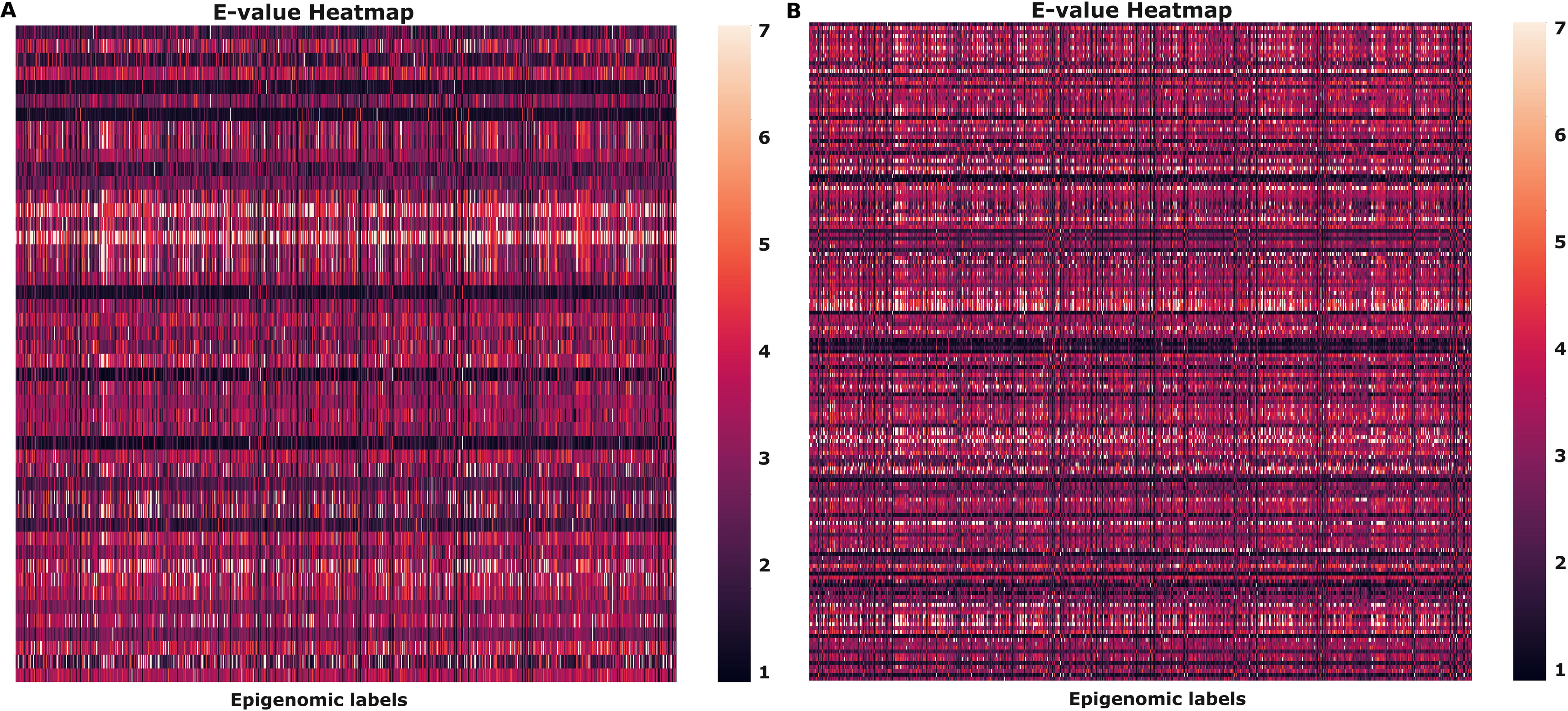
E-value heatmap. Heatmap showing significant variants A. Neurological condition specific GWAS SNPs, B. Neurological eQTL SNPs. Variants in rows and labels in columns.

The significant gene-variant pair files [v8] of the following samples belonging to the nervous system from GTEx portal were used for extracting neurological condition specific eQTL positive variant set: “Cerebellum”, “Nucleus accumbens basal ganglia”, “Cortex”, “Caudate basal ganglia”, “Cerebellar Hemisphere”, “Anterior cingulate cortex BA24”, “Amygdala”, “Spinal cord cervical C-1”, “Hypothalamus”, “Substantia nigra”, “Frontal Cortex BA9”, “Hippocampus”, “Putamen basal ganglia”. Top 1000 hits from each sample were used for the predictions and after stringent filtering, 169 highly significant regulatory SNPs were predicted by the model [Figure 4B, Supplementary Table S3].

### ASD GWAS variants

The variant effect predictions were restricted to 92 ASD SNPs from GWAS Catalog and 4 highly significant regulatory ones were predicted by the model [Figure 5]. The significant labels and their E-values are shown in Supplementary Table S4.

**Figure 5:**
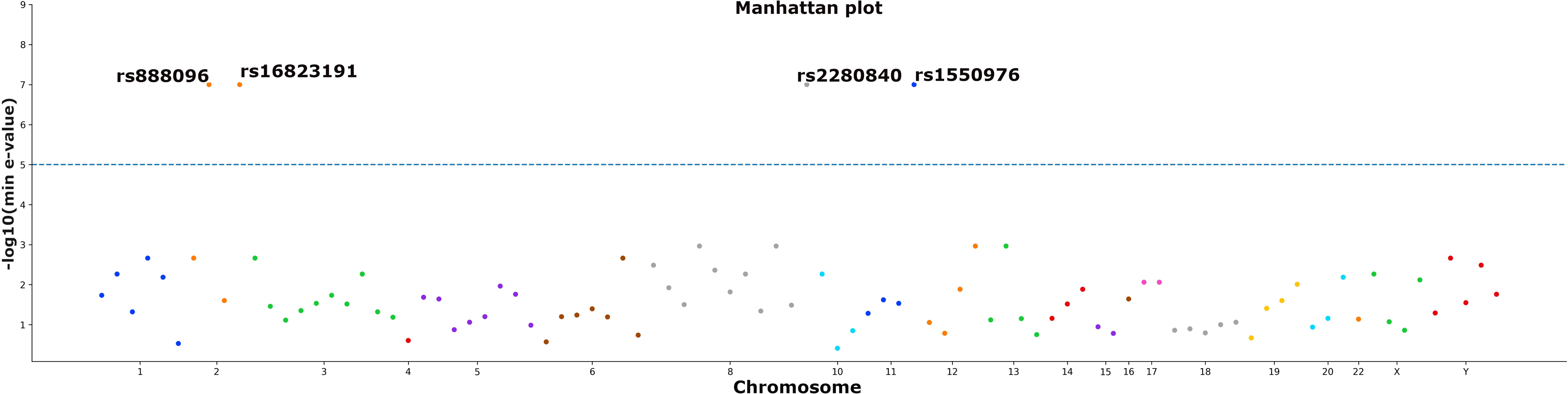
Manhattan plot of ASD regulatory variants predicted by Neur-Ally. ASD associated SNPs from GWAS Catalog having significant E-values are shown above the horizontal threshold line.

### Regulatory brain variants

The model was tested for variant effect prediction on reported probable regulatory variants in the brain. 8 such SNPs were selected from publications [23–29] and their differential epigenomic changes upon in silico mutagenesis were predicted by the model [Table 1]. Predictions of differential epigenomic labels for rs7364180 and rs12411216 were found to be significant. rs7364180 is found to be associated with many eQTL genes in brain tissues [28]. rs12411216 is reported to be a probable regulatory risk variant for mild cognitive impairment in Parkinson’s disease [29].

**Table 1:** Neur-Ally predictions for reported probable brain regulatory variants. . a) rsIDs of the regulatory SNPs, b) Functional effect reported in the publications and their reference number in superscript, c) Minimum E-value predicted by the model, d) Epigenomic label having the lowest E-value.

| **SNP<sup>a</sup>** | **Functional effect<sup>b</sup>** | **Min.**<br>**E-value<sup>c</sup>** | **Min E-value Label<sup>d</sup>** |
| --- | --- | --- | --- |
| rs405509 | Decreased APOE expression in brain tissue<sup>(17)</sup> | 0.0115 | Layer of hippocampus (H3K27me3) |
| rs2526377 | Increased expression of PICALM and TGFBR1 in neuronal progenitor cells<sup>(18)</sup> | 0.016 | Astrocyte (H4K20me1) |
| rs6733839 | BIN1 depletion in microglia<sup>(19)</sup> | 0.0649 | Motor neuron (H3K27ac) |
| rs1990620 | Frontotemporal dementia risk, increases CTCF recruitment<sup>(20)</sup> | 0.0005 | Dorsolateral prefrontal cortex (chromatin accessibility) |
| rs356168 | Parkinson's disease risk, regulates SNCA gene expression<sup>(21)</sup> | 0.004 | SK-N-SH (ZC3H10 ChIP-Seq) |
| rs1476679 | Alzheimer's disease risk, associated with eQTL genes<sup>(22)</sup> | 0.0005 | SK-N-SH (POLR2AphosphoS5 ChIP-Seq) |
| rs7364180 | Alzheimer's disease risk, associated with eQTL genes<sup>(22)</sup> | 0.0 | Dorsolateral prefrontal cortex (CTCF binding) |
| rs12411216 | Parkinson's disease risk, regulates glucocerebrosidase expression<sup>(23)</sup> | 0.0 | Dorsolateral prefrontal cortex (chromatin accessibility) |

For the SNP rs7364180, the model predicted significant chromatin accessibility changes in samples like motor neuron, head of caudate nucleus, brain microvascular endothelial cell and cell lines like SK-N-SH, H54, Daoy and A172. In addition to that, significant changes were also found in TF ChiP-Seq labels from dorsolateral prefrontal cortex, PFSK-1 and SK-N-SH. Neur-Ally also predicted significant changes in chromatin accessibility because of rs12411216 in dorsolateral prefrontal cortex and posterior cingulate gyrus.

## DISCUSSION

Quantitative genetic studies like GWAS and eQTL analysis in neurological disorders have revealed several risk variants in the non-coding regions of the genome. The functional consequences of the coding variants can be interpreted by the effect of the mutation on the protein structure. But for non-coding variants, the regulatory consequence can be specific to the cell or tissue type. Experimental methods to determine the effect of all the significant variants from a study will be difficult because of this heterogeneous nature. Computational tools developed from the publicly available epigenomic datasets can be used to create prediction models for this purpose. Such functional predictions will help to differentiate between actual causative variants and those which are highly linked to them.

In this study, we have created a deep learning model named Neur-Ally, trained it on epigenomic datasets derived from nervous tissues and cells. Most of the existing variant effect prediction models were trained on regulatory datasets from multiple tissues or cell line samples. It will be difficult for such models to learn the regulatory signatures that are cell or tissue specific. In case of diseases where changes in a particular organ or tissue contribute majorly to the pathophysiology, models trained on that particular tissue or cells will be more helpful. Machine learning model trained on human retinal epigenomic datasets was developed to predict the effect of non-coding variants in human retinal cis-regulatory elements [30]. Another pancreatic islets specific model trained on multiple epigenome profiling datasets was used for prioritizing type 2 diabetes association signals [31]. Machine learning model for variant effect prediction in Alzheimer’s disease was developed and achieved better accuracy compared to other models [32]. The model was trained on 39 features of which 9 were regulatory ones. Data from seven brain tissues were used to form regulatory regions. In contrast, Neur-Ally was developed and trained on multiple brain samples including tissues, cells, in vitro differentiated cells, primary cells and organoids. 758 regulatory datasets were used to create a training dataset of 9 million genomic bins [200bp] overlapping epigenetic features. Hence Neur-Ally is important in the sense that it is a variant prediction model that uses a large number of neuronal specific regulatory datasets and can be used for all neuronal disorders. The model predicts the regulatory labels upon giving nucleotide sequences of genomic bins as input. Model achieved commendable performance while predicting TF binding, Histone modifications and Chromatin accessibility upon testing. In silico mutagenesis was carried out after training the model and the significant regulatory effects of neurological condition specific mutations was identified. Regulatory consequences were identified in neurological condition specific GWAS, eQTL, ASD GWAS and reported probable regulatory neurological condition specific variants.

Immune system abnormalities are identified in patient categories with neurological disorders. Thus, the associated genetic variants may have regulatory functions in non-neuronal samples as well. The data pre-processing scripts available with Neur-Ally can be used for creating training datasets with analysis files of different sample types. For the neuronal predictions, additional epigenomic data can be added whenever they are available. Therefore, the prediction performance of the model that we have developed can further be improvised as and when the newer datasets arrive.

## DATA AVAILABILITY

Neur-Ally is open source and available at [https://github.com/ anilprakash94/neur_ally]. The epigenomic datasets are available at https://www.encodeproject.org/

## CONTRIBUTIONS

A.P. and M.B. conceptualized the work, A.P. and M.B. performed the analysis, A.P. and M.B. interpreted and wrote the manuscript.

## ACKNOWLEDGEMENTS

We are grateful to the Dept. of Biotechnology, Govt. of India for providing the intramural support to Rajiv Gandhi Centre for Biotechnology to MB, and Council of Scientific and Industrial Research [CSIR] for Senior Research Fellowship to A.P.

## SUPPLEMENTARY INFORMATION

**Supplementary Table S1:** Metric values for individual epigenomic labels. The AUROC and PR_AUC values are given in separate columns and their mean values are provided at the end of the file.

**Supplementary Table S2:** GWAS SNPs of neurological disorders with significant regulatory predictions. The “Signif. labels” column contains the epigenomic labels in which significant regulatory changes were predicted and the respective E-values are given in the “Signif. scores” column.

**Supplementary Table S3:** Top brain eQTL SNPs with significant regulatory predictions. The “Signif. labels” column contains the epigenomic labels in which significant regulatory changes were predicted and the respective E-values are given in the “Signif. scores” column.

**Supplementary Table S4:** Autism Spectrum Disorder GWAS SNPs with significant regulatory predictions. The “Signif. labels” column contains the epigenomic labels in which significant regulatory changes were predicted and the respective E-values are given in the “Signif. scores” column.

